# Computational modeling reveals biological mechanisms underlying the whisker-flick EEG

**DOI:** 10.1101/2024.12.13.628364

**Authors:** Joseph Tharayil, James B. Isbister, Esra Neufeld, Michael Reimann

**Affiliations:** Blue Brain Project, École polytechnique fédérale de Lausanne (EPFL) Campus Biotech, Geneva, Switzerland; IT’IS Foundation, Zurich, Switzerland

## Abstract

Whisker flick stimulation is a commonly used protocol to investigate somatosensory processing in rodents. Neural activity in the brain evoked by whisker flicks produces a characteristic EEG waveform recorded at the skull, known as a somatosensory evoked potential. In this paper, we use *in silico* modeling to identify the neural populations that serve as sources and targets of the synaptic currents contributing to this signal (presynaptic and postsynaptic populations, respectively). The initial positive deflection of the EEG waveform is driven largely by direct thalamic inputs to Layer 2/3 and Layer 5 pyramidal cells, though interestingly, L5-L5 inhibition plays a modulatory role, reducing the amplitude and width of the deflection. This suggests that increasing thalamocortical connectivity and decreasing L5-L5 inhibition may be responsible for some of the changes observed in the EEG waveform over the course of development. The negative deflection is driven by a more complex mix of sources, including both thalamic and recurrent cortical connectivity. We demonstrate that small changes to the local connectivity of the circuit, particularly to perisomatic inhibitory targeting, can have an important impact on the recorded EEG, without substantially affecting firing rates, suggesting that EEG may be useful in constraining *in silico* neural models.

## 1 Introduction

The somatosensory evoked potential (SEP) is the EEG signal evoked by somatosensory stimuli, recorded at the skull. SEPs are commonly studied in the context of the whisker flick paradigm in rodents for the study of sensory processing, integration, and plasticity [1]. Understanding the whisker flick SEP can provide insight into the biophysical basis of somatosensory processing and the related EEG. Following a whisker flick, the triggered SEP has a stereotypical waveform with stimulus intensity-dependent amplitude, consisting of a positive deflection (P1) with a width of around 5 ms, immediately followed by a negative deflection (N1) of the same width [2]. It is believed [2] that the rise of the P1 component is driven by activation of thalamocortical synapses, but the cause of its decay is not fully understood. In contrast to the view that P1 is fully thalamocortically driven, the P1 component does not manifest in the EEG signal until around postnatal day 13, suggesting that the signal arises due to activity in supragranular layers that appears with maturation of local connectivity [3], and cannot be explained solely by thalamic input. However, the local connectivity patterns that permit the emergence of the P1 component are not known.

Experimental studies [2] have shown that blocking inhibitory synaptic activity alters the shape of the late phase of the N1 component, but does not affect the P1 component. However, the degree to which this effect can be attributed directly to decreased inhibitory post-synaptic currents in pyramidal neurons, as opposed to broader changes in circuit-level activity, is unclear. Modulation of the GABAA receptor has been shown to cause changes to the voltage-sensitive dye (VSD) signal that depend at least in part on changes in disynaptic inhibition [4]. Computational modeling at the circuit level may be very useful in understanding the impact of inhibition on the N1 component of the SEP.

The Blue Brain Project (BBP) model of the rat non-barrel primary somatosensory cortex (nbS1) consists of ∼ 4.2 million reconstructed neurons with accurate morphologies, optimized physiological properties, and algorithmically generated connectivity [5, 6]. In this study, we simulate a subvolume of the BBP nbS1 model with *∼* 210,000 neurons (Fig. 1a), which has been shown to produce realistic firing activities in response to a simulated whisker-flick stimulus modeled by activating virtual thalamic fibers that innervate the cortical model (Fig. 1b-d, [6]). The peri-stimulus-time histograms (PSTHs) of firing in layer-wise excitatory/inhibitory populations closely match *in vivo* data on a millisecond timescale [6], thus implying a high-level of precision in the population responses studied here. To simulate the EEG signals associated with our whisker flick stimulus simulation, we use a detailed finite element electromagnetic model of the rat head in conjunction with BlueRecording [7], a set of software tools, which enables the simulation of extracellular recordings from neural circuit models by calculating a “weights file” that specifies the contribution weights of transmembrane currents from each compartment in the model to the signal.

**Figure 1:**
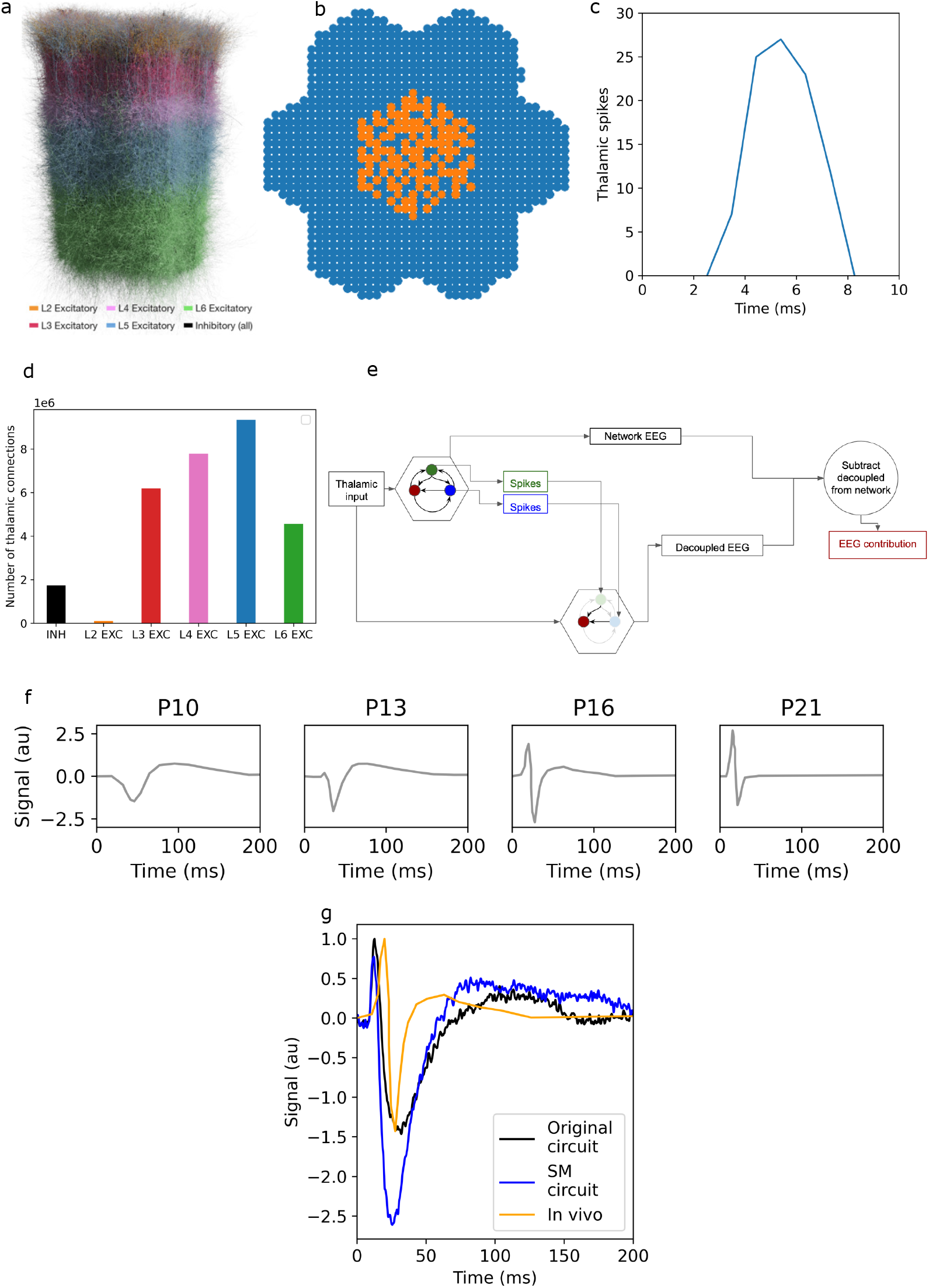
a: Seven column subvolume of nbS1 used in this study. b: Whisker flick is simulated by activating a subset of thalamic fibers innervating the cortex. Activated fibers are marked in orange. Fibers not activated are marked in blue. c: Thalamic spike times are drawn from an *in vivo* peri-stimulus-time histogram (PSTH) recorded in response to whisker flick [13] d: Thalamic innervation of nbS1 primarily targets excitatory populations. e: Cartoon of the workflow used to isolate the contribution of a specified presynaptic population to the SEP. Thalamic spikes are played into a circuit (hexagon) with several interacting populations (for clarity, only three are shown). The EEG calculated from the circuit is referred to as the “network EEG”. To calculate the contribution to the EEG from the population represented by the red circle, the spikes from the other two populations are recorded, and played into a circuit in which only synaptic connections to the population represented by red circle are active. The EEG recorded from this simulation is referred to as the “decoupled EEG”. The decoupled EEG is subtracted from the network EEG to obtain the contribution of the population represented by the red circle. f: *In vivo* SEPs recorded at various ages from postnatal day 10 (P10) to postnatal day 21 (P21), digitized from [3] and normalized to the peak P1 amplitude on postnatal day 16. g: Comparison of SEP signals from the original and SM circuits with *in vivo* data obtained on postnatal day 16 [3]. The *in vivo* signal and the *in silico* signal from the original circuit are normalized to their respective P1 peaks; the signal from the SM circuit is normalized to the P1 peak from the original circuit.

In particular, we aim to identify the contributions of different presynaptic and postsynaptic populations to the SEP. Here, the contribution of a presynaptic population to the SEP refers to the direct impact of the synaptic currents generated by the presynaptic population on the SEP; we do not include the downstream (i.e., multi-synaptic) effects of these currents. Similarly, the contribution of a postsynaptic population to the SEP refers to the direct impact of all of the transmembrane currents in that population on the SEP. While disambiguating the contributions of specific postsynaptic populations would be effectively impossible *in vivo*, doing so is trivial *in silico*. BlueRecording reports the contribution of each neuron in the model to the SEP; these neuronal contributions are summed in postprocessing to generate population contributions [7]. In this paper, we report the contributions of postsynaptic cells in Layers 2/3, 4, 5, and 6; this is effectively equivalent to the contribution of excitatory postsynaptic cells in the corresponding layers, as the contributions of inhibitory cells are negligible (Supplementary Figure S.2)

Identifying the contribution of presynaptic populations to the SEP relies on a feature of our simulation tools referred to as “spike replay” [8]. This feature allows us to provide a list of neuron IDs and corresponding spike times. During the simulation, the efferent synapses of the provided neurons will be activated at the prescribed times, without affecting the behavior of the neurons themselves. Moreover, spikes fired by these neurons as a result of network interactions during the simulation do not additionally activate any synapses. To isolate the contribution of an individual presynaptic population of interest to the recorded SEP, we add all neurons in the model to this list except for the presynaptic population of interest, and ensure that no spikes are transmitted from the population of interest. Subtracting the results of this simulation from those of an original simulation with no spikes removed yields the contribution to the SEP of the population in question (Fig. 1e).

One cannot isolate the impacts of particular cortico-cortical pathways by simply setting connection weights between the populations of interest to zero, and subtracting the resulting SEP from the original signal, as this would have unpredictable effects on the firing rates of the postsynaptic population, and consequently on downstream populations. It would therefore be impossible to disambiguate the effects of the removed synaptic inputs from the broader changes in circuit activity, which may well deviate unrealistically from *in vivo* conditions. The problem has been previously approached with a similar solution to ours, namely replaying only the spikes from the presynaptic population of interest [9], but we believe that our approach is more informative because the approach in [9] removes the majority of spikes, which could potentially have significant effect on the membrane dynamics of the postsynaptic neurons (which in normal conditions are highly leaky), thus affecting the SEP.

In principle, it is possible that even our procedure of disconnecting the circuit and playing in the majority of the spikes from the connected simulation could lead to an entirely different dynamical regime. If this were the case, the procedure would not produce accurate estimates of the contributions of the presynaptic populations. However, as the sum of the contributions of all presynaptic populations approximates the original signal (Fig. S.1), it is unlikely that the procedure places the network in a very different dynamical regime.

The approach we implement in this paper yields the contributions of synapses from an identified presynaptic population.However, that contribution can vary depending on the locations of synapses with respect to postsynaptic morphologies. For example, [10] predicts strong unitary local field potentials of inhibitory spikes in hippocampus due to their perisomatic locations. To explore the impact of synapse location on the EEG signals produced in our simulations, we calculate the SEP from a rewired version of the somatosensory cortex model [11], referred to as the Schneider-Mizell (SM) circuit, which incorporates more accurate local connectivity obtained from the MiCrONS electron microscopy dataset [12].

We compare the SEPs generated *in silico* to *in vivo* results from [3] (Fig. 1f). In particular, we are interested in how our model compares to *in vivo* data recorded on postnatal day 16 (P16), as our model is based largely on data obtained from an animal on P14, with additional data from older animals. Because our model encompasses only a small subset of the somatosensory cortex, and because we do not apply the amplification used in [3], we cannot expect the amplitude of the *in silico* SEP to match that of the *in vivo* signal. We therefore normalize both the *in vivo* signal and the *in silico* signals from the original circuit to their respective P1 peaks, comparing only the shape of the signals. The *in silico* signal from the Schneider-Mizell circuit is normalized to the P1 peak of the original circuit.

## 2 Results

### 2.1 SEPs modeled *In silico* approximate *in vivo* recordings

We compare the SEPs produced by our circuit models to *in vivo* SEPs in P16 [3]. We note that the *in vivo* recordings were conducted with a reference electrode over the lambda point, while *in silico* recordings were conducted with a reference over the hindlimb region. However, we show that changing the reference location only affects the SEP by a linear scaling factor (Supplementary Fig. S.3).

The peak amplitude of the P1 component of the *in vivo* SEP on P16 is a factor 104 greater than that of the *in silico* SEP from the original circuit. Our model thus produces EEG signals with amplitude on the same order of magnitude as the (unamplified) signals recorded in [3], where an amplifier with gain 5000X is used.

Given that our model does not encompass the full brain, and therefore should be expected to produce EEG amplitudes somewhat smaller than *in vivo*, the fact that it produces EEG signals that are approximately 50% the *in vivo* amplitude is an important validation for our model.

We then find that the relative N1 amplitude of the *in vivo* SEP also matches that of the original *in silico* SEP (Fig. 1g). However, the P1 component *in vivo* occurs approximately 5 ms later than *in silico*, and the relative amplitude of the N1 component of the SEP from the SM circuit is substantially greater than *in vivo* (Fig. 1g). The width of the N1 component *in silico* is roughly three times that of the *in vivo* SEP, though the SM circuit produces an N1 component 25% narrower than the original.

Our results indicate that the model successfully replicates the key features of the SEP. However, the duration of the simulated P1 component is a better match to the *in vivo* data than that of the N1 component. The observation that the width of the N1 is narrower in the SM circuit than in the original, while the amplitude is larger in the SM circuit, suggests that this rewiring has mixed effects on the realism of our model.

### 2.2 The P1 component is mostly driven by thalamic input

The isolated thalamic contribution to the SEP is quite similar to the SEP obtained from the original nbS1 (Fig. 2 a.i). However, the isolated thalamic contribution produces a slightly larger P1 component. The difference in the P1 component can be attributed to the smaller postsynaptic contribution from L2/3 and, particularly, L5 in the circuit, than what would be evoked by the thalamic contribution alone (Fig. 2 a.ii and a.iv). We also note that the postsynaptic contribution from Layer 4 in the connected circuit has a sharp negative peak during the P1 component (Fig. 2a.iii); however, the impact thereof on the overall signal is limited, as the postsynaptic Layer 4 contribution is an order of magnitude smaller than that of other layers (see Supplementary Materials Section A.4).

**Figure 2:**
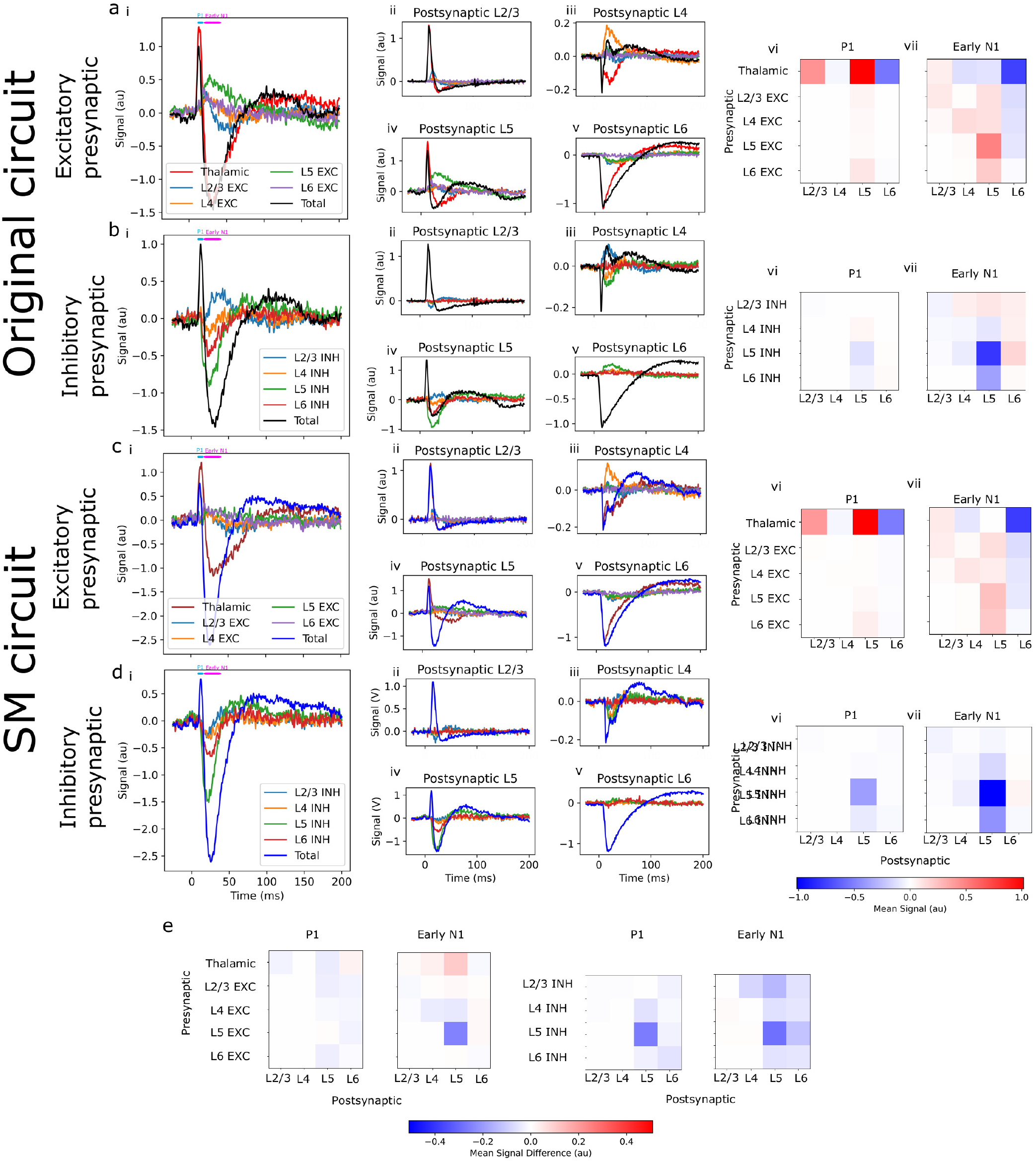
a.i: SEP from the original nbS1, and the contributions to the SEP of the different excitatory presy-naptic populations. Timing of the P1 component (defined as the period from 8-16 ms post-stimulus) and the early N1 component (defined as the period from 16-40 ms post-stimulus) are marked with blue and pink bars, respectively. a.ii-v: Postsynaptic contributions from the different layers, attributable to excitatory presynaptic input from corresponding populations. a.vi-vii: Mean contributions to the SEP over the P1 component and early N1 component, by excitatory presynaptic and postsynaptic population. b: Same as panel a, but for inhibitory presynaptic populations. c: Same as panel a, but for the SM nbS1. d: Same as b, but for the SM nbS1. e: Differences between the SM and the original circuit in the mean postsynaptic contribution from the different layers, attributable to presynaptic input from corresponding populations.

In addition to the original nbS1 model, we also simulate the SM circuit. In the SM circuit, PV+ interneurons target perisomatic regions with more specificity than in the original circuit (see [5] for details). As in the original circuit, the P1 component in the SM circuit is smaller than what would be evoked by the thalamic contribution alone (Fig. 2c.i), due primarily to differences in the postsynaptic Layer 5 contribution (Fig. 2c.iv).

The difference between the total signal and the contribution of thalamic input to the signal is evidently the contribution of recurrent connectivity within the circuit. We further split this contribution into the individual contributions from pre- and post-synaptic populations (see Methods). Contributions from Layer 1 are not shown, as they are very small. (See Supplementary Fig. S.1 and Supplementary Materials Section A.1 for why the sum of the presynaptic contributions does not add up perfectly to the total signal).

For both the original circuit (Fig. 2b.iv, b.vi) and the SM circuit (2d.iv,d.vi), the reduction in the amplitude of the P1 component can be attributed to the impact of L5-L5 inhibition. The greater reduction in amplitude in the SM circuit can be attributed to a stronger impact of L5-L5 inhibition (Fig. 2e). This suggests that reduction of this inhibitory connectivity may be partially responsible for the increased amplitude of the P1 component over the course of development (Fig. 1f, see also Discussion). However, as recurrent activity only changes the amplitude of the P1 component by *∼* 15%, the primary contributor to P1 is clearly the thalamic input.

### 2.3 L5-L5 inhibition has a significant impact on N1

In the original circuit, the N1 component is similar to that produced by the thalamic input alone (Fig. 2a.i). Unlike for the P1 component, however, this does not result from a dominating thalamic contribution to the N1. While thalamic inputs to Layer 6 substantially contribute to the N1 (Fig. 2a.v, a.vii), and in particular prescribe the time course of the recovery, thalamic inputs do not dominate as they do in the case of the P1 component. We observe that during the early part of the N1 component (prior to *∼* 40 ms post-stimulus), inhibitory input from L5 has a strong driving effect on the contribution of L5 to the N1 component, as does, to a lesser extent, inhibition from L6 to L5 (Fig. 2b.iv,b.vii). However, this is partially compensated for by the effect of cortico-cortical excitation, particularly L5-L5 (Fig. 2a.iv,a.vii), which has the opposite effect on the early N1.

Compared to the original circuit, the SM circuit has a significantly stronger N1 component (Fig. 2c,d), due to a stronger negative contribution from postsynaptic L5 cells (Fig. 2d.iv). Presynaptically, this is explained by a weaker positive contribution of L5 excitation to the early N1, and by a stronger negative contribution of L5 inhibition (Fig. 2e). As for the original circuit, the early N1 component in the SM circuit is driven primarily by L5-L5 inhibition, with a smaller role for L6-L5 inhibition (Fig. 2d.i,d.iv,d.vii).

We observe that the postsynaptic contribution of Layer 4 during the early N1 component is strongly affected by recurrent connectivity, both in the original circuit (Fig. 2a.iii) and in the SM circuit (Fig. 2c.iii). In both cases, this difference can be attributed primarily to the effect of L4-L4 excitation (Fig. 2a.vii,c.vii). However, this has a much smaller effect on the N1 component than recurrent connectivity in Layer 5, since the magnitude of the postsynaptic Layer 4 contribution to the EEG is an order of magnitude smaller than that of Layer 5 (see Supplementary Materials Section A.4).

That the SEP is so strongly affected by recurrent inhibition suggests that maturation of inhibitory circuit connectivity, particularly within L5, may be responsible for changes in the N1 component observed during development.

### 2.4 Perisomatic targeting explains the difference between the original and the SM circuit

We found that the contribution of L5-L5 excitation to the SEP is greater in the original than in the SM circuit (Fig. 2e, 3a.i). While firing rates in Layer 5 pyramidal cells do have a narrower peak and longer tail in the SM circuit than the original (Fig. 3a.ii), it is unclear if this difference is large enough to explain the difference in the contribution to the SEP. The difference in the contribution of L5-L5 excitation to the SEP could potentially be attributed to changes in membrane excitability due to rewiring of inhibitory connections.

**Figure 3:**
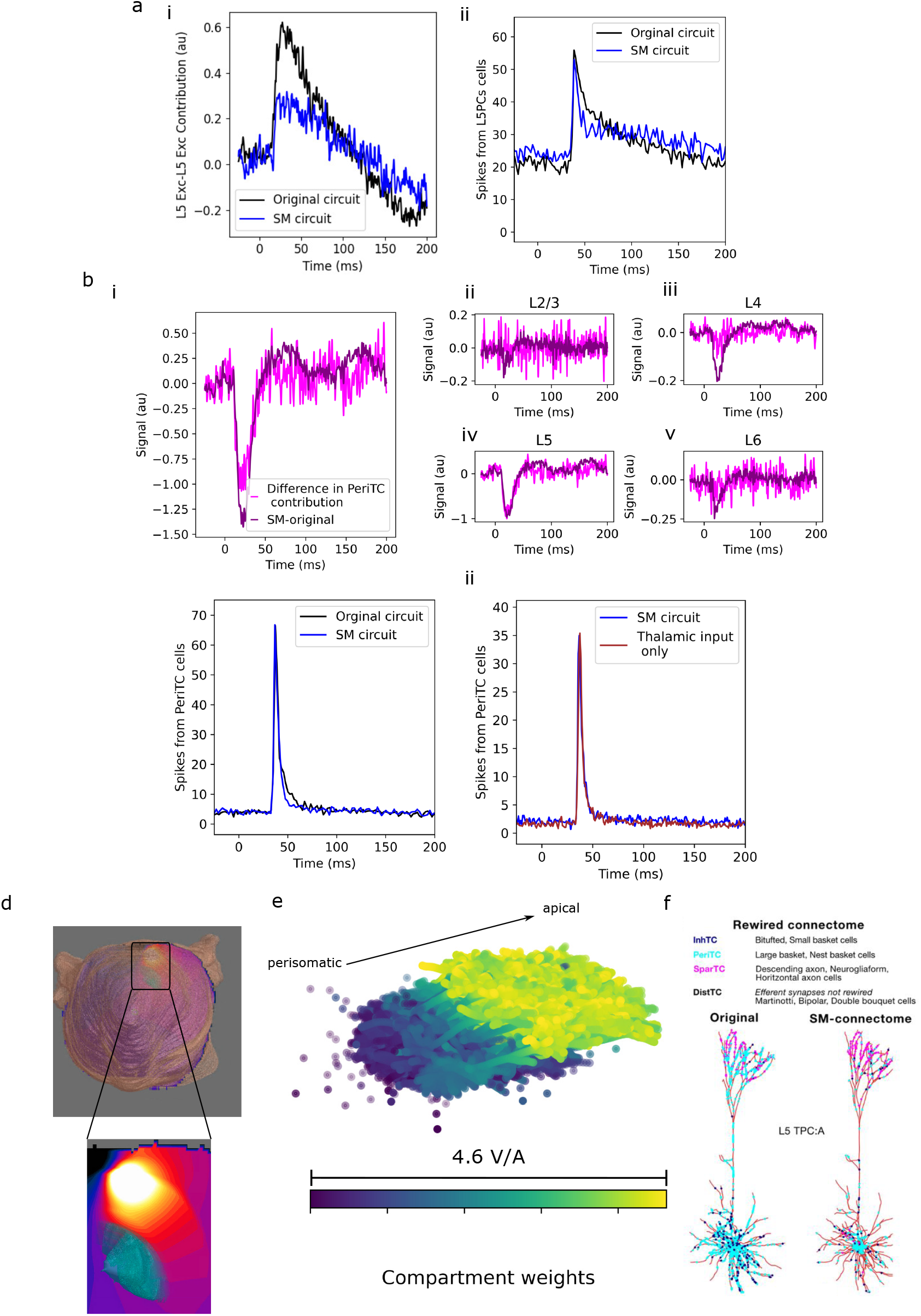
a: i. Contribution of L5-L5 exitation to the SEP in the original and SM circuits. ii: Firing rates of Layer 5 pyramidal cells in the original and SM circuits. b: i. The difference in contribution to the SEP due to presynaptic excitatory input from L5 perisomatic targeting cells accounts for the difference between the SM and the original circuit. ii-v: Differences in the postsynaptic contribution from each layer, attributable to L5 PeriTC presynaptic input. Note the difference in y-axis scales between panels. c; i: Firing rates for Layer 5 PeriTC cells do not differ substantially between the original and the SM circuit. ii: Firing rates for Layer 5 PeriTC cells do not differ substantially between the SM and the disconnected circuit. d: Potential induced by current applied between recording electrodes in a finite element model of the rat head. Skin visualized in beige. Zoom: Potential over the somatosensory cortex (visualized in light blue) e: Sensitivity of the EEG signal to currents over the membrane of Layer 5 pyramidal cell morphologies. Each electrical compartment of a neuron is indicated as a dot at its center; the sensitivity is reflected in its color. f: Rewiring moves PeriTC synapses on pyramidal cells to perisomatic compartments (reproduced from [6] with permission; “SM-connectome” indicates the SM circuit)

The difference between the SM and the original circuit can be primarily attributed to the presynaptic contribution of Layer 5 large basket cells (LBCs) and Layer 5 nest basket cells (NBCs) (collectively called perisomatic targeting cells (PeriTCs)) to Layer 5 pyramidal cells (Fig. 3.b). We note, however, that while the difference in the PeriTC contributions between the SM and original circuits is close to the total difference, the contributions of populations, such as thalamic inputs to L5, and inhibitory inputs from other layers to L5 and L6, are also substantial (Fig. 2e), but largely cancel out.

For PeriTCs, synapse locations are closer to the soma in the SM circuit. The firing rates of PeriTCs do not differ significantly between the original and the SM circuits (Fig. 3c.i). Moreover, the firing rates of PeriTCs do not differ substantially between the SM circuit and a circuit with disconnected cortico-cortical connectivity that receives only thalamocortical input (Fig. 3c.ii), implying that the activity of these cells is driven by direct thalamic input (as expected, given that inhibitory cells respond to thalamic stimulus with low latency [14]). This indicates that the difference in their contributions must be attributable to the difference in their synapse locations, which demonstrates that changes in local connectivity can have relevant impacts on EEG signals, without significantly affecting their firing rates. This confirms Rimehaug et al., 2023 [15] who found a similar dissociation between firing rates and electrical signal, but for the current source density (CSD) signal instead of the EEG.

In the SM circuit, PeriTC synapses are moved from apical compartments to perisomatic compartments (Fig. 3f). For a random sample of L5PCs in our model, we visualize the sensitivity of the EEG to transmembrane currents in each compartment (see [7] for details). Note that because of the gauge degree-of-freedom of the electric potential, these “compartment weights” are only defined up to a constant offset, i.e., only the gradient in weights over a neuron affects the signal. We therefore refer to “more negative” and “more positive” weights to mean weights closer to one end or the other of the range. We observe that the perisomatic compartments have more negative weights than the apical compartments (Fig. 3e). With synapses primarily targeting perisomatic regions, the neuron acts as a dipole, with currents at the soma and associated return currents along the apical dendrite. Because of the large gradient in weights between the soma and apical dendrites, concentrated inhibitory synaptic input near the soma would lead to a more negative deflection in the EEG than the same input distributed over the dendritic arbor.

### 2.5 Specificity of layer 5 inhibition drives differences in width of the N1 circuit

As previously described, the width of the N1 component *in silico* is approximately 3 times longer than that of the *in vivo* signal (Fig. 4a). The SM circuit produces an N1 component 25% narrower than the original, with a full width at half maximum (FWHM) of 21 ms and 28 ms respectively. We denote the width of the N1 component in the original circuit *FWHM*_*o*_ and that in the SM circuit *FWHM*_*r*_. We denote the difference in the width of the N1 component between the original and SM circuits as Δ*FWHM* .

**Figure 4:**
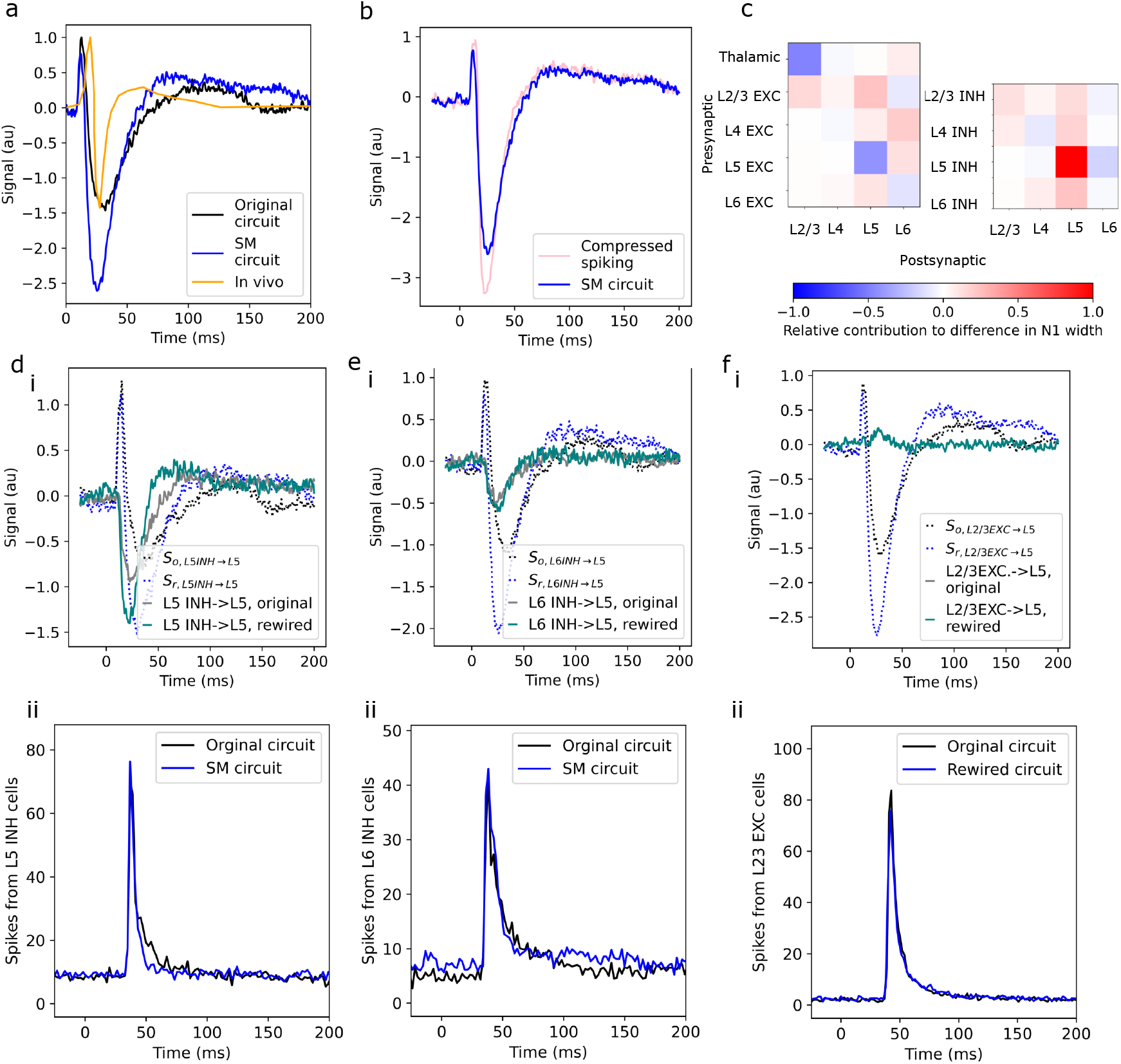
a: Comparison of SEP signals from the original and the SM circuit with *in vivo* data obtained on postnatal day 16 [3]. b: Compressing the timing of spikes from Layer 5 PeriTC neurons does not have a substantial impact on the SEP. c: Contributions of different presynaptic-postsynaptic pathways to the difference in the width of the N1 component between the original and the SM circuit. A positive contribution implies that the pathway narrows the N1 in the SM circuit. d: i. Contribution of L5 - L5 inhibition to the SEP, for both the original and SM circuits (solid lines). SEP from the two circuits without the contribution of L5 - L5 inhibition (dashed lines). ii: Firing rate of L5 inhibitory cells in the original and SM circuits. e, f: Same as d, but for L6 inhibitory cells and L2/3 excitatory cells, respectively.

In order to determine whether more precise spike timing in the PeriTC cells would bring the SEP from the SM circuit closer to *in vivo* observations, we replay the spikes from the fully connected circuit into a disconnected circuit, with all PeriTC spikes occurring between 10 ms post-stimulus and 40 ms post-stimulus shifted to 15 ms post-stimulus (i.e., the peak of the spike histogram (Fig. 3b.i)). Even though this level of compression is highly exaggerated, the resulting SEP is not substantially different from the SEP in the SM circuit – the amplitude of the N1 component is slightly greater, and the time course of the N1 is slightly faster (Fig. 4b).

In order to determine the source of the reduction in the width of the N1 component, we calculate, for each combination of presynaptic population *x* and postsynaptic population *y*, the SEP signal with the contribution of that particular presynaptic-postsynaptic pathway removed, in both the original and the SM circuit. We denote the resulting signals *S*_*o*,*x→y*_ and *S*_*r*,*x→y*_ for the original and the SM *™* circuits, respectively. The width of the N1 component in each of these signals is *FWHM*_*o*,*x→y*_ and *FWHM*_*r*,*x→y*_. We denote the difference *FWHM*_*o*,*x→y*_ − *FWHM*_*r*,*x→y*_ = Δ*FWHM*_*x→y*_.

The extent to which a particular presynaptic-postsynaptic pathway contributes to the difference Δ*FWHM* in the width of the N1 component in the SM circuit relative to the original circuit can be inferred by the effect of removing the pathway. Thus, for each pathway, we calculate Δ_*x→y*_ = (Δ*FWHM −* Δ*FWHM*_*x→y*_)*/*Δ*FWHM*. A positive value of Δ_*x→y*_ implies that removing the pathway *x → y* reduces the difference Δ*FWHM* between the N1 component of the original and SM circuits, and therefore, that the pathway *x → y* could contribute to the observed difference.

We find that L5-L5 inhibition contributes strongly to Δ*FWHM* (Fig. 4 c.ii), as do, to a lesser extent, L5-L5 excitation (Fig. 4 c.ii), and L2/3 excitation to L5 (Fig. 4 c.i). The time constant of recovery of the contribution of L5-L5 inhibition to the SEP is faster in the SM circuit than in the original (Fig. 4d.i). Due to the impact of noise on the signal when removing populations (see Supplementary Materials Section A.1), we were unable to attribute the narrowing of the N1 component to a more specific population. However, as firing rates for Layer 5 excitatory cells are not substantially different between the original and SM circuits (Fig. 4d.ii), the differences in the time course of the SEP contribution must be due to differences in synapse placement in the SM circuit. Unlike the contribution of L5-L5 inhibition to the SEP, the contributions of L6-L5 inhibition and L2/3-L5 excitation do not differ substantially between the original and SM circuits (Fig. 4e.i, f). These populations contribute to the increased width of N1 in the original circuit only because they are a relatively larger share of the N1 component in the original circuit than in the SM circuit.

## 3 Discussion

We have simulated the somatosensory evoked potential induced by a whisker flick in two *in silico* models of the rat non-barrel primary somatosensory cortex. In both cases, the resulting SEP has the basic shape observed *in vivo*, with an initial positive deflection (P1) followed by a negative deflection of similar amplitude (N1). Through careful simulation techniques we were able to decipher the contributions of individual neuronal populations to the characteristics of the two components.

We have shown that the main features of the SEP, namely the N1 and the P1 component, can in principle be generated entirely by direct thalamocortical input, with the N1 component being driven by thalamic input to L6 neurons (Fig. 2a.vii). The P1 component is primarily driven by direct thalamic input to L2/3 and L5, but it is, to a certain extent, modulated by L5-L5 inhibition (Fig. 2b.iv,b.vi, d.iv,d.vi). While it is not surprising that P1 is primarily thalamocortically driven, the modulatory role of L5-L5 inhibition has not, to our knowledge, been previously reported.

*In vivo*, the P1 component of the whisker flick EEG emerges around postnatal day 13 and increases in amplitude over the course of development [3]. Our finding that the P1 component is driven primarily by thalamic input indicates that the emergence of the P1 component over the course of development is not primarily related to changes in local connectivity, though decreases in L5-L5 inhibition during maturation may partially account for the observed increase in the amplitude of the P1 component. Rather, we speculate that the increased P1 component may result from increased thalamic input to L2/3 and L5, or morphological changes in L2/3 and L5 pyramidal cells. It has previously been shown that the strength of thalamic innervation of L5 pyramidal cells does increase over the course of development [16]. Further *in silico* and *in vivo* experiments could test these hypotheses.

While thalamic input to L6 contributes to the N1 component – particularly to its late phase – recurrent connectivity has a larger impact on N1 than thalamic innervation. In particular, cortico-cortical inhibition has a substantial impact on the amplitude of the N1 component (Fig. 2b.vii, d.vii), though in the original circuit, this is balanced by the effect of cortico-cortical excitation (Fig. 2a.i,a.iv,a.vii). While it is unsurprising that overall cortico-cortical inhibition plays a significant role in the N1 component, our simulations have been able to reveal the specific pre- and post-synaptic populations that contribute to the N1.

Our results contrast sharply with those of Bruyns-Haylett et al. [2], who found that the application of a GABA antagonist had no effect on P1 or the initiation of N1, but increased the width of the N1. In contrast, we find that inhibition plays a significant role in the initiation of N1, while the width of N1 is determined primarily by thalamic input to Layer 6 (Fig. 2). It is important to note, however, that the experiments in [2] used adult animals, and moreover the intervention in that study was applied in a “closed-loop” manner; i.e., the application of the GABA antagonist could have effected the spiking activity of local circuits, which was explicitly avoided in our approach. While [2] found no significant differences in resting-state MUA after the application of the GABA antagonist, it is unclear what effects it had on evoked activity. Given the complexity of the contributions to the N1 component in our model, it is difficult to predict how a GABA antagonist would effect the EEG in a closed-loop format; future work that explicitly models the GABA antagonist in a closed-loop configuration may shed help explain the findings in [2].

The contribution of a pre- and post-synaptic neuron population to the signal depends on – among other factors – the locations of the synapses between the populations. By comparing contributions for two circuits with different locations (“original” vs. “SM”), we were able to improve out understanding of this. Our main finding was that increasing specificity of inhibitory connectivity in the form of targeted innervation by Basket Cells around Pyramidal Cell somata led to a slight decrease of the P1 amplitude and a significant increase of the N1 amplitude, combined with a narrowing of its width. These changes are not explained by changes in pre-synaptic spiking, but in how the synaptic currents associated with the spiking affect the signal. Our result confirms a similar dissociation between spiking and the CSD signal described before [15].

*In vivo*, the N1 component is present from at least postnatal day 7. Its width peaks on postnatal day 10, before narrowing as the animal ages; after P16, its amplitude begins to decrease slightly [3]. In our model, the ratio of the amplitude of the P1 and N1 components is similar to that on P16 in the original circuit, and between that of P13 and P16 in the SM circuit. The N1 component has a width closest to, but somewhat larger than, the width of the *in vivo* signal on P10.

The width of the S1 component in the SM circuit more closely resembles that of the *in vivo* signal than that of the original circuit. We have shown that this is largely due to the impact of L5-L5 inhibition. It has been found that activation of distal synapses can lead to longer-lasting depolarization [17]; the reduction in N1 width in our SM circuit may therefore be attributable to the more specific somatic targeting of inhibition.

Still, the duration of the N1 component in our models is longer than *in vivo* by a factor of *∼* 3. The time course of the N1 component is largely driven by thalamic input to Layer 6 (Fig. 2a.v,c.v), which dominates the late N1. It may be that our model overestimates the time constant of the thalamic contribution to the L6 SEP. Thalamic innervation in our model is determined by the depth profile of thalamic synapses over the cortical column, without accounting for cell-type or neurite-type specific targeting [5]. It is therefore possible that – if thalamic fibers preferentially target L5 cells, or basal dendrites – our model overestimates thalamic input to L6. In fact, previous studies have found a unimodal distribution of thalamic synapses onto L5 pyramidal cells, primarily targeting basal dendrites [18]; if this preference holds for all cell types, it may be that many of the thalamocortical synapses onto aprical dendrites of L6 pyramidal cells in our model should be moved to basal dendrites of L5 pyramidal cells. As thalamic input to L5 also produces a slow negative contribution to late N1 (Fig. 2a.iv, c.iv), this would not entirely account for the increased duration of the N1 in our model. However, as discussed previously, activation of distal synapses leads to a longer-lasting depolarization than activation of proximal synapses [17]. Thus, moving thalamocortical synapses from L6 apical dendrites to L5 basal dendrites may result in a faster negative deflection in the thalamic → L5 contribution to the SEP than observed in our present models.

Alternatively, it is possible that the short N1 component observed *in vivo* emerges through cancellation of the late part of N1 by other populations. As discussed above, L5-L5 inhibition can shorten the N1 component in the rewired circuit; it is possible that our model does not capture similar effects from other populations, potentially those from regions outside of the simulated subvolume. It has been suggested that, beginning around 25 ms post-stimulus, activity in the contralateral hemisphere has a significant effect on the recorded SEP [19]; this aligns with the time course of the termination of the N1 component *in vivo*. Activation of the contralateral hemisphere increases over the course of development, correlating with a reduction in the width of the N1 component [3]. Given that our model produces an N1 component with width close to that of the *in vivo* N1 on P10, before the contralateral hemisphere begins to be activated, it is possible that inter-hemispheric activity, which is not represented in our model, is responsible for the reduction in the width of the N1.

We observed that the onset of the SEP in our model occurs earlier than *in vivo*. This may be attributed to differences in the timing of thalamic spikes in our model compared to *in vivo*. While spike times in our model are based on an *in vivo* PSTH [13], this data was obtained from adult animals, while the *in vivo* SEPs were obtained from juveniles. As the onset of the SEP becomes earlier as the animal matures [3], it seems likely that this explains the discrepancy in the timing of the SEP. In our model, axonal delays for the transmission of thalamic action potentials are calculated based on the distance from the bottom of Layer 6 to the synapse location, ignoring the length of the axon in the thalamus itself; this may also contribute to the faster latency in our model.

We were surprised by the fact that the incorporation of connectivity trends from the MiCrONS dataset worsened the agreement in peak-to-peak amplitude between our model and the *in vivo* data. The *in vivo* SEPs presented in this paper were taken from an animal on postnatal day 16, and the *in silico* model was built using data from a variety of sources with varying ages. The MiCrONS dataset is, in contrast, obtained from a mature animal [12]. It may be that the connectivity trends observed there emerge later in development. Because maturation is associated with a reduction, rather than increase, in the amplitude and width of the N1 component [3], this alone does not account for our observations. It may be that our *in silico* model also excludes other changes in anatomy and physiology over the course of maturation which compensate for the deepening of N1 by increased specificity of PeriTC targeting.

Our approach to deciphering the contributions of populations relies on the assumption that the contributions of pre-synaptic populations sum up approximately linearly. This assumption has been used and validated before for the current source density (CSD) signal [9]. Additionally, we provide validation for the EEG signal in this work. However, we also found that rewiring inhibitory connectivity modulated how excitatory pre-synaptic populations contributed to the signal even though their spiking activity remained unchanged. This demonstrates complex, non-linear interactions between populations and the limitations of the assumption. While this assumption may be approximately true for a given circuit, it must be re-evaluated *globally* even for small *local* changes to a model.

### 3.1 Limitations

A potential limitation of this study is the application of whisker-flick stimulus to non-barrel somatosensory cortex; it is not clear that nbS1 would produce the same responses to thalamic input as barrel cortex. However, this is not a fatal limitation, as the thalamic input is modeled on innervation of the barrel cortex [5] and the model has been shown to replicate neural activity in the barrel cortex in response to whisker flick and optogenetic manipulation [6]. It has been shown to reproduce both the population-level magnitude and time course of spiking responses to these stimuli.

While our *in silico* models are extensively validated, there remain aspects that may differ from *in vivo* conditions in ways which might influence the calculated SEP. The simulated synaptic physiologies have been shown to produce excitatory post-synaptic potentials with *in vivo*-like amplitude and time course, but this data has been recorded at the soma, rather than in the dendrites. Similarly, the electrical models of the neurons have been validated on the basis of responses to stimuli recorded at the soma [6]. Particularly for pyramidal cells, with their extended dendritic arbors, the contribution to the EEG is strongly influenced by transmembrane currents in the dendrites as well as the soma. Thus, differences in the electrophysiological properties of the dendrites may influence the simulated SEP.

We observed that relatively small alterations to local connectivity may significantly impact the simulated SEP. While the large scale connectivity trends in our circuit model are well validated [5], local connectivity may benefit from the incorporation of more data obtained from electron microscopy or other methods. In particular, it may be necessary to incorporate connectivity data obtained at specific points during maturation. The observation that our model replicates the width of the *in vivo* P1 component but not the N1 component, coupled with the observation that the P1 component is driven primarily by thalamic input, while the N1 component depends more strongly on recurrent inhibition, may suggest that our model better matches the thalamocortical connectivity of the P16 animal used in [3] than its recurrent connectivity.

### 3.2 Future directions

This work suggests that the increase in the amplitude of the P1 component observed over the course of development may be due to increased thalamocortical connectivity to L2/3 and L5 pyramidal cells. Reducing the strength of L5-L5 inhibition may be responsible for the reduction in N1 amplitude observed over development, though as previosuly discussed, inputs from outside the somatosensory cortex are also likely to contirbute to the N1. Further research is needed in order to fully explain the changes in the SEP over the course of development. It is almost certain that changes in cellular anatomy and physiology during the course of development also impact the SEP. To a limited extent, the influence of changes in anatomy on the SEP can be approximated by scaling the weighting factors for each neural compartment to mimic the effects of a spatial rescaling of the neuron. However, it may be necessary to generate new morphologies and reoptimize neuron electrical models to fully capture the cellular-level changes that occur during development.

We have shown that changes in local connectivity can cause significant changes in EEG, without strongly affecting firing rates (Fig. 3). This suggests that simulated EEG signals may be useful as a secondary metric, in addition to firing rates, to constrain *in silico* models. The use of simulated electrical signals, in the form of current source densities (CSD), has previously been proposed as a metric to constrain model building [15]. Indeed, validating the CSDs produced by our model, may be useful in constraining modifications made to replicate EEG in different states and stages of development.

## 4 Acknowledgments

This work was supported by funding to the Blue Brain Project, a research center of the École polytechnique fédérale de Lausanne (EPFL), from the Swiss government’s ETH Board of the Swiss Federal Institutes of Technology.

## 5 Author contributions

Conceptualization: J.T., E.N. and M.W.R. Methodology: J.T. and M.W.R. Software: J.T. Validation: J.B.I. Formal Analysis: J.T. and J.B.I. Investigation: J.T. Resources: E.N. and M.W.R. Writing-Original Draft: J.T. Writing: Review and Editing: J.T., J.B.I., E.N. and M.W.R. Visualization: J.T. Supervision: E.N. and M.W.R.

## 7 Resource Availability

### 7.1 Lead Contact

Requests for further information should be directed to Joseph Tharayil.

### 7.2 Materials availability

This computational study used no materials.

### 7.3 Code availability

The simulations performed in this study are run using the Neurodamus [8] simulation control tool, which integrates with the CORENEURON [20] computation engine. To calculate EEG signals, Neurodamus relies on a “weights file”, which lists the sensitivities of the EEG to the currents from each neural compartment in the model. Weights files for the calculation of EEG signals are created using BlueRecording [7].

- Code for running the simulations and generating the figures in this paper is available at https://github.com/joseph-tharayil/whiskerFlick. Postprocessed EEG traces from these simulations are available on Zenodo under the following DOI: `10.5281/zenodo.14442089`.
- The source code for BlueRecording is available at https://github.com/BlueBrain/BlueRecording.
- The seven column subvolume of the BBP circuit model is available under the following DOI: `10.5281/ zenodo.7930276`. The SM circuit is available under the following DOI: `10.5281/zenodo.10677883`.
- Membrane mechanism files used in the neuro-simulations are available at https://github.com/BlueBrain/neurodamus-models.
- Neurodamus is available at https://github.com/BlueBrain/neurodamus.
- CoreNEURON itself is fully integrated into the NEURON simulation environment, which is available at github.com/neuronsimulator/nrn.
- Code for the generation of FEM models, as well as a list of dependencies, are available at https://github.com/BlueBrain/BlueBrainHeadModels. The required finite element meshes are available on Zenodo (DOI: `10.5281/zenodo.10926947`).
- Finite element meshes and FEM output files used in the simulations are available on Zenodo (DOI: `10. 5281/zenodo.10927050` and oSparc (https://osparc.io/#/study/2d25d5be-b667-11ef-baa3-0242ac177740).

## 8 Methods

### Neural circuit models

We simulate the central, 7-column subvolume of the BBP model of the nbS1 [6]. This subvolume contains *∼* 210,000 biophysically-detailed neurons, which receive Ornstein-Uhlenbeck conductance noise from virtual sources, to represent the effect of synapses from non-modelled regions. Noise input is optimized to produce *in silico* firing rates equal to desired ratios (*P*_*FR*_) of *in vivo* firing rates, corresponding to different levels of excitability (in this paper, *P*_*FR*_ = 0.3), with the ratio of the standard deviation of the noise to the mean of the noise fixed at 0.4 (*R*_*OU*_ = 0.4)[6]. Simulations are conducted with a simulated extracellular calcium concentration (which modulates the efficacy of synaptic transmission) of 1.05 mM. The subvolume is innervated by virtual thalamic fibers representing VPM inputs; a virtual whisker flick stimulus activates 10% of VPM fibers, with randomly-sampled spike times obtained from an *in vivo* PSTH of VPM activity in response to whisker deflection, as in [6]. The delay between thalamic spike times and synapse activation (accounting for axonal transmission time) is based on the distance from the bottom of Layer 6 to the synapse location.

In addition to the original nbS1 subvolume, we also study a rewired circuit [11], in which the targeting of interneurons is modified to more closely match the MiCrONS dataset [12, 5]. We refer to this circuit as the Schneider-Mizell (SM) circuit. Parvalbumin-positive (PV+) interneurons target perisomatic regions with more specificity, vasoactive intestinal peptide-expressing (VIP+) interneurons preferentially target inhibitory neurons, and Layer 1 and neurogliaform interneurons have primarily monosynaptic connections. Synapses and input noise parameters are recalibrated for the SM circuit, in order to ensure that firing rates match *in vivo* data [6].

### Calculating EEG signals

We calculate the EEG signal from our neural circuit models using the “reciprocity approach” [21]: The contribution to the EEG of a transmembrane current in a particular neural compartment is the product of the current magnitude and the electric potential that would be produced at the location of the compartment by a unit current applied between the recording and the reference electrode. The workflow for calculating EEG signals using BlueRecording is described in detail in [7] and briefly summarized below:

- We created a finite element model of the rat head, with a recording electrode directly over the forelimb region of the somatosensory cortex and a reference electrode over the hindlimb region. Using Sim4Life (Zurich MedTech AG, Zurich, CH), we calculated the electric potential in the head generated by a current applied between the electrodes.
- Using BlueRecording, we interpolated the potential field at the location of each neural compartment in the model and wrote the values to a “weights file”.
- At runtime, the Neurodamus [8] simulation control program loads the weights file and, at each time step, calculates the EEG as the dot product of the transmembrane currents and the weights. At the end of the simulation, an HDF5 file is produced, which reports the contribution to the EEG for each neuron.
- In postprocessing, we summed the contributions of all neurons from populations of interest.

### SEP simulations

#### Unmodified circuits

For both the original and the SM circuit, we simulate 10 trials, each of which has a different random seed for the conductance noise and the selection of thalamic fibers. We refer to this set of trials as a *simulation campaign*. In each trial, two thalamic stimuli are simulated. The EEG signal is calculated as described above; in postprocessing, we isolate the SEP produced by the second whisker-flick stimulus, in order to account for the effects of accommodation. We apply a second-order bandpass Butterworth filter with cutoff frequencies of 1 Hz and 500 Hz. The SEP is averaged over the 10 trials. Signals from both the original and SM circuits are normalized to the peak amplitude of the P1 component in the *in silico* EEG from the original circuit.

#### Isolating postsynaptic contributions

BlueRecording outputs the contribution to the EEG signal from each neuron individually. Because of the linearity of Maxwell’s equations, we can isolate the contribution of a particular postsynaptic population by summing the contributions of all neurons belonging to that population. In this paper, we isolate the postsynaptic contributions of inhibitory neurons in Layer 1, and both excitatory and inhibitory neurons in layers 2/3, 4, 5, and 6.

#### Isolating presynaptic contributions

To isolate the contributions of specific presynaptic populations, we begin with the spike output files from each simulation in the simulation campaign on the fully connected circuit. Because the spike output files associate each spike in the simulation with the unique ID of the spiking neuron, we can filter out the spikes from the population of interest to create a series of “spike input files” (one per simulation in the original campaign). We then create a new simulation campaign, in which we set synaptic transmission triggered by spikes fired during the simulation to 0. Pre-recorded spikes that are replayed into the circuit are not affected by this manipulation. In addition to the thalamic input, the “spike input files” are replayed into the circuit. We ensure that the outcomes of stochastic processes, such as synaptic release and stochastic ion channels, are preserved between the simulations from the fully connected circuit and those from the disconnected circuit. The SEPs are calculated for the new campaign as described above, and subtracted from the SEPs from the fully-connected campaign. This difference represents the contribution of the presynaptic population to the SEP (Fig. 1e).

To isolate the contributions of thalamic populations, we replay the unaltered spikes from the fully-connected simulations into a disconnected circuit, without the thalamic input, and subtract the resulting EEG from that of the fully-connected simulation.

In this paper, we isolate the presynaptic contributions of inhibitory neurons in Layer 1, and both excitatory and inhibitory neurons in layers 2/3, 4, 5, and 6, as well as the VPM inputs.

### Comparison with *in vivo* data

To compare the SEPs generated *in silico* with *in vivo* data, we digitized the highest-amplitude SEP recorded on postnatal day 16 in [3], as the highest-amplitude SEPs were recorded above the somatosensory cortex. The *in vivo* EEG is normalized to its P1 peak.

---

**Figure S.1:**
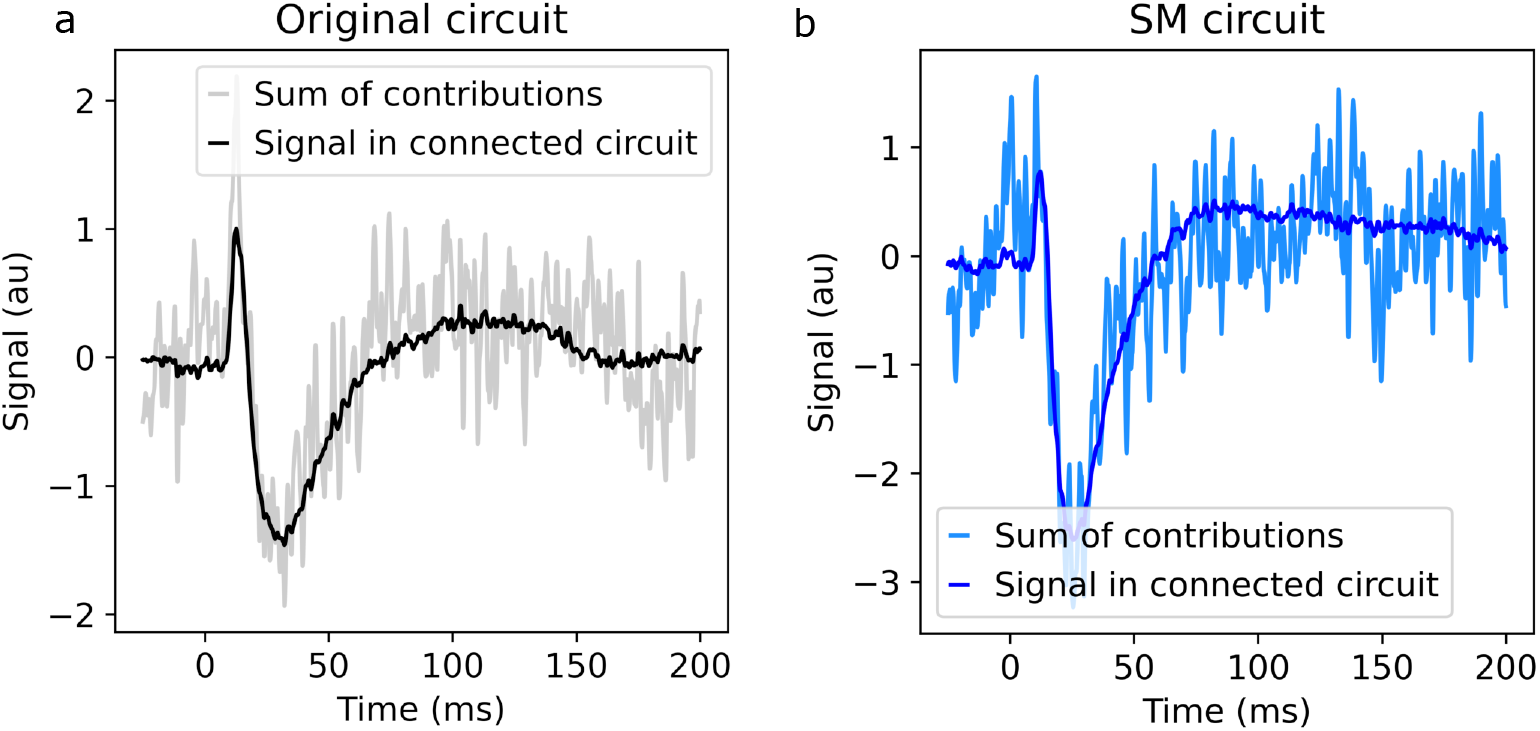
a: SEP from the original circuit, compared to the sum of the calculated contributions from each cortical population and the thalamic input. b: SEP from the SM circuit, compared to the sum of the calculated contributions from each cortical population and the thalamic input.

## A Supplementary material

### A.1 Sum of estimated presynaptic contributions approximates total signal

To validate the approach of using spike replay to isolate the contributions of individual cortical populations to the SEP, we compare the SEP generated by the fully-connected circuit to the sum of the contributions of each presynaptic neural population, as defined in Sec. 8 (including the cortical populations and the contribution of the thalamic input). For both the original and the SM circuits, the sum of the population contributions does approximate the original signal (Supplementary Fig. S.1). However, much more high-frequency noise is present in the reconstructed signal than in the original signal. This can be explained by differences in spike timing between the fully-connected and the disconnected simulations, which is in turn a consequence of the “missing” spikes in the input replayed into the disconnected simulation (see Sec. 8). When the EEG from the disconnected simulation is subtracted from that of the fully-connected simulations to calculate the contribution of the presynaptic population, high-frequency noise results from the differences in the timing of the action potentials’ contribution to the signal; this noise is amplified as the contributions of the presynaptic populations are summed.

### A.2 Postsynaptic inhibitory cells do not contribute to EEG

The EEG signal is dominated by excitatory postsynaptic cells, in both the original and the SM circuits (Supplementary Fig. S.2). This is expected, as a cell’s contribution to the EEG is driven by the difference in the compartment contribution weights between locations of input currents and return currents; this difference is small for interneurons.

### A.3 Reference electrode location does not affect signal shape

We demonstrate that the choice of reference electrode location (S1HL or lambda), has little impact on the shape of the SEP (Supplementary Fig. S.3). All electrodes are modeled as spheres. For simulations with the S1HL reference, the electrodes have a radius of 0.1 mm; for simulations with the lambda reference, the electrodes have a radius of 0.5 mm.

### A.4 Postsynaptic contributions from L4 are small compared to other layers

We observed that the contribution of postsynaptic Layer 4 cells to the EEG is an order of magnitude smaller than that of other layers (Fig. 2). Factors influencing the magnitude of the contribution from a postsynaptic population include the number of cells in the population, the correlation in the contribution of individual cells, and the magnitude of the contribution from each cell. The magnitude of the cellular contribution is in turn influenced by the range of the compartment weights over the cell (a larger range of weights leads to a larger contribution), the alignment of synaptic and return currents with the axis over which the weights vary, and the amplitude of the currents themselves.

For a random sample of neurons in each population, we calculate the average range in compartment weights over the neuron (difference between 90th percentile and 10th percentile). The range of weights of Layer 5 cells are, on average, more than twice that of L2/3 and L4 cells (Supplementary Table A.4). However, the L2/3 population is larger than L4 and L5. Were these factors to scale proportionally, we would expect L2/3 and L4 to have similar contribution magnitudes, Band that of L5 to be *∼*2 times larger. The observed contribution from Layer 4 is therefore smaller than expected. The whisker flick stimulus effectively synchronizes activity between neurons [6], which would tend to increase the amplitude of the EEG contribution. Therefore, we speculate that this difference can instead be attributed to a lower amplitude of current arriving in L4, or to these currents not aligning with the axis of the weights over L4.

**Figure S.2:**
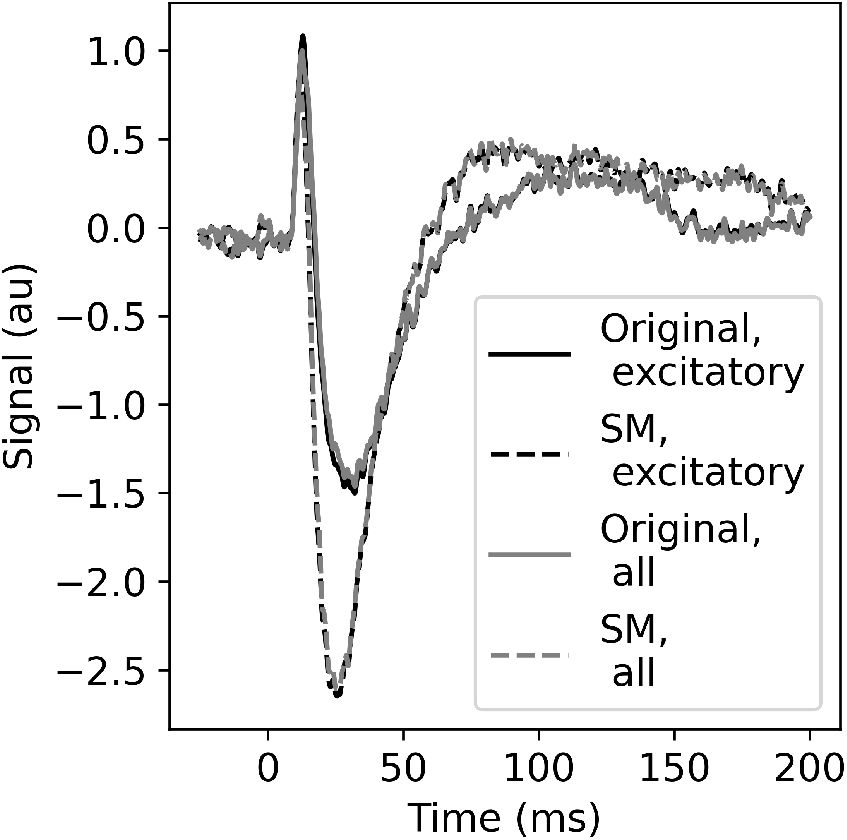
The contribution to the *in silico* EEG from excitatory postsynaptic cells alone is almost identical to the total EEG, in both the original and the SM circuits.

**Figure S.3:**
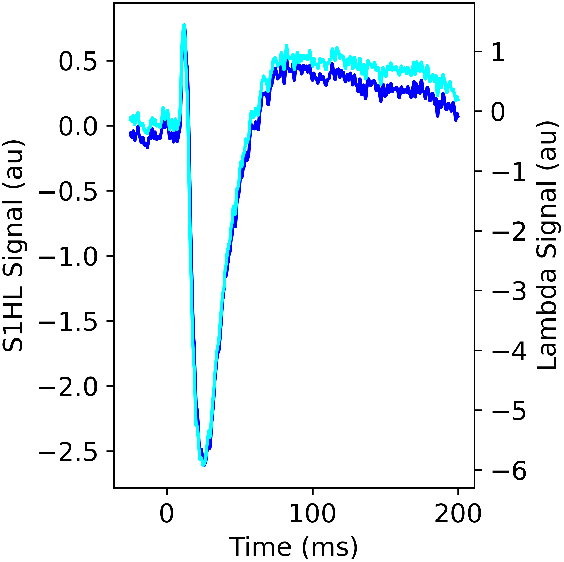
SEP recorded in the SM circuit from an EEG electrode over the S1FL region, with the reference electrode over the S1HL region or the lambda point.

**Table 1:** Factors influencing postysnaptic contributions for selected populations.

| Table 1: Factors influencing postsynaptic contributions for selected populations |  |  |  |
| --- | --- | --- | --- |
| Population | Cell count | Weights range (V/nA) | Peak-to-peak SEP Amplitude (V) |
| L2/3 | 53000 | $1.2 \times 10^{-9}$ | $4.7 \times 10^{-8}$ |
| L4 | 29000 | $1.5 \times 10^{-9}$ | $8.6 \times 10^{-9}$ |
| L5 | 35000 | $3.1 \times 10^{-9}$ | $7.6 \times 10^{-8}$ |

